# Enhanced hypoxic ventilatory response is accompanied by a decrease in carotid body mitochondrial oxygen affinity in heart failure

**DOI:** 10.64898/2026.11.06.764874

**Authors:** Agnieszka Swiderska, Mohammed Obeidat, George W.P. Madders, Alice S. Whitley, Jagen C. Burke, Menglu Lin, Gina L. J. Galli, Andrew W. Trafford

## Abstract

Carotid bodies (CBs), the main peripheral oxygen sensors, play a significant role in sympathetic overdrive in heart failure (HF), resulting in worse clinical outcomes. Whilst CBs are an attractive therapeutic target, the development of CB focused therapies has been limited as the oxygen sensing mechanism is not fully understood. It has been hypothesised that the oxygen sensor is part of mitochondrial complex IV, and reactive oxygen species (ROS) signalling plays a crucial role in the oxygen sensing cascade. However, changes to CB mitochondrial function have never been determined in HF. Hence, we aimed to address this gap by characterising CB mitochondrial function and ROS production using a rapid pacing ovine model of HF. The hypoxic ventilatory response (HVR) and autonomic function were assessed pre- and post HF induction. Tissue homogenates were prepared from whole CBs from control and HF and utilised to measure kinetics of mitochondrial oxygen affinity, respiration and H_2_O_2_ production rates by high-resolution respirometry. In HF, HVR and sympathetic drive were increased, while complex IV oxygen affinity was decreased. Despite an increase in abundance of mitochondrial complexes, no changes in mitochondrial respiration or H_2_O_2_ production were observed. However, aconitase activity and lipid peroxidation level were increased in HF. Overall, these results suggest that the CB mitochondrial function is altered in HF, and decreased CB mitochondrial oxygen affinity may suggest a higher complex IV oxygen sensitivity. Higher oxygen sensitivity accompanied by an increased oxidative stress may be key drivers of CB overactivity and therefore autonomic imbalance in HF.

## Introduction

The carotid body (CB) is the main peripheral sensor of oxygen. A fall in partial pressure of arterial oxygen (P_a_O_2_)^1^ increases CB output which increases sympathetic activity and evokes cardiorespiratory changes, namely an increase in minute ventilation (V_E_) and cardiac output^2^. While this system is vital for physiological homeostasis, the CB can become dysfunctional in pathological conditions, such as heart failure (HF), hypertension, cardiometabolic diseases leading to higher sympathetic drive and worse clinical outcomes^3–8^.

HF is characterised by reduced cardiac contractility resulting from dysregulated Ca²⁺ handling, alongside reduced catecholamine responsiveness and compensatory sympathetic overactivation^9^. Chronic sympathetic overactivation can exacerbate cardiac dysfunction and accelerate disease progression^10^. Current HF therapies include β-blockers which are focused on normalising β-adrenergic signalling to lower sympathetic drive; however, their effectiveness remains limited and mortality rates remain high^11^. Increased sympathetic outflow is driven at least partially by CB overactivation, and critically, enhanced CB function is associated with lower survival rates in HF patients and poorer quality of life^12^.

Furthermore, bilateral CB resection has been shown to have beneficial effects both in pre-clinical trials and patients ^13–15^.

In agreement with these epidemiological findings, CB chemoreflex sensitivity has been widely reported to be increased in preclinical small animal models and HF patients^12,15–20^. Early studies provided evidence that under normoxic (21% FiO_2_) conditions, HF rabbits had an increased carotid sinus nerve activity, V_E_ and renal sympathetic nerve activity^17,18^.

Furthermore, the magnitude of the V_E_ increase in response to hypoxia (10 – 15 % FiO_2_) was greater in the HF group, therefore suggesting an enhanced CB sensitivity to hypoxia in these animals^17,18^. While CB resection was considered as a promising new therapeutic avenue, a follow-up study performed on HF patients who underwent bilateral CB resection 5 years prior, reported potential adverse effects, including nighttime oxygen desaturation and greater variability in oxygen saturation during exposure to mild hypoxia (15% FiO_2_)^13,21^.

Consequently, CB resection as a treatment for HF should be approached with caution. Alternative therapeutic strategies that aim to normalise, rather than completely abolish, chemoreflex activity should therefore be considered^21^. However, realising this potential requires a greater understanding of the mechanisms underpinning CB dysfunction in HF.

The prevailing theory of oxygen sensing is that the CB type I cell have unique mitochondria with specialised complex IV subunits having unusually low apparent oxygen affinity and therefore can sense minute changes in arterial oxygen level^2^. The mitochondria-to-membrane hypothesis states that in response to hypoxia, complex IV is inhibited which slows the flow of electrons and ultimately results in reverse electron transport (RET) back to complex I and increase in ROS production^22,23^. ROS then signal to the cell membrane to trigger neurotransmitter release, carotid sinus nerve activation and the central cardiorespiratory response to hypoxia. Type I cell sensitivity to hypoxia is increased in HF and is accompanied by an increase in angiotensin II signalling, NADPH oxidase 2 activity and augmented oxidative stress (both cytosolic and mitochondrial; for review see^24^). However, because these studies were performed on isolated CB type I cells, the contribution of mitochondrial remodelling to CB hypoxia sensitivity remains unknown.^25,26^ This study aimed to fully characterise the effects of HF on CB mitochondrial function, including oxygen affinity, oxidative capacity and ROS production. We hypothesised that oxygen affinity would be lower in the HF, whilst ROS production would be increased. Using a sheep tachypacing HF model, we observed a decrease in oxygen affinity of CB complex IV, which may contribute to increased CB hypoxic sensitivity in HF. CB mitochondrial oxidative capacity was decreased in HF, mainly due to an increase in LEAK respiration. Surprisingly, the abundance of mitochondrial complexes was increased, but not their activity. At the same time, there was a decrease in mitochondrial oxidative stress (potentially due to mild uncoupling of mitochondria), while the whole tissue oxidative stress was elevated which could be driving increased CB activity and ultimately lead to sympathetic overdrive in HF.

## Methods

### Animals

All animal experimentation was performed in accordance with The Animals (Scientific Procedures) Act 1986 and EU Directive 2010/63. Local ethical approval was obtained from The University of Manchester Animal Welfare and Ethical Review Board. The reporting of use of animals follows the ARRIVE II principles^27^. All procedures were performed using female Welsh Mountain sheep (37 animals, ∼ 18 - 24 months of age, 36.8 ± 0.9 kg). Animals were fed *ad libitum* hay supplemented with ruminant concentrate with free access to drinking water and maintained at ∼ 21°C on a 12:12 hour light:dark cycle. No animals were excluded from the study or post data acquisition analysis.

### Induction of Heart Failure

Animals were randomly assigned to control and HF groups. HF was induced by right ventricular tachypacing (210 beats per minute) as described in detail previously ^28–31^. Briefly, under general anaesthesia (2.5 – 3% isoflurane in oxygen) IS1 pacemaker leads were inserted via the jugular vein and positioned in the right ventricular apex and connected to a dual chamber pacemaker located subcutaneously in the pre-scapular region. Perioperative analgesia (Metacam, 0.5 mg.kg^-1^, subcutaneous) and antibiosis (Betamox LA, 15 mg.kg^-1^, intramuscular) were provided. Animals were allowed to recover from the procedure for at least 7 days post-surgery before tachypacing was commenced. Animals were monitored at least once day for subjective signs of HF (lethargy, breathlessness).

### Hypoxic ventilatory response

A subgroup of animals (n = 9) underwent spirometric analysis under general anaesthesia (2.5 – 3.5% isoflurane in oxygen). Animals were intubated and breathed spontaneously. Baseline breathing (100% FiO_2_ with 2.5 - 3.5 % isoflurane, later referred to as Nx) was recorded for approximately 10 minutes after which animals were exposed to hypoxia (11.5% FiO_2_ balanced with nitrogen) for up to 5 minutes or until the blood oxygen saturation level measured by pulse oximetry fell to 75%. Animals were allowed to recover from the hypoxic challenge before recovery from anaesthesia or euthanasia (if spirometry was performed at terminal HF stage). The peak hypoxic response was analysed from a 20 second window of stable baseline breathing. The change in minute ventilation (ΔV_E_) was calculated by subtracting the baseline V_E_ from peak hypoxic response V_E_ and was used as method to evaluate the change in CB hypoxic sensitivity with HF.

### *In vivo* monitoring

Sheep underwent weekly conscious transthoracic echocardiography to assess cardiac function (described in detail in ^32^).

A subgroup of sheep (n = 7) also underwent *in vivo* assessment of autonomic nervous system function ^30,33,34^. Briefly, electrocardiogram (ECG) recordings were performed to measure heart rate and following, a baseline ECG recording, the β_1_ adrenoceptor blocker metoprolol (0.5 mg.kg^-1^; metoprolol tartrate made up in saline solution) was used to assess sympathetic input. The ECG was recorded for ∼ 3 minutes following injection and the final 20 seconds of the recording after injection were used for analysis.

### Carotid body dissection

Sheep were injected intravenously with 20,000 IU heparin to prevent tissue blood coagulation prior to euthanasia with an overdose of pentobarbital sodium (Pentoject; 200 mg.kg^-1^). An incision was made along the jugular groove to expose the common carotid artery and its bifurcation. A large area surrounding the common carotid artery was dissected out. The CBs were then rapidly identified and dissected out. Samples were either taken for homogenate preparation or snap frozen and stored in liquid nitrogen for subsequent molecular analyses. CBs were randomly assigned to each experiment, with no specific side (left or right) being taken for a specific experiment.

### Homogenate preparation

Following dissection, a single CB was placed in an ice-cold Mir05 buffer containing (in mmol.L^-1^): sucrose, 110; K-MES, 60; 4-(2-hydroxyethyl)-1-piperazineethanesulfonic acid, 20; monopotassium phosphate, 10; magnesium chloride, 3; ethylene glycol-bis(ß-aminoethyl ether)-N,N,N’,N’-tetraacetic aicd, 0.5; and 0.1% bovine serum albumin (BSA). Tissue was homogenised using a hand-held homogeniser. Homogenate solution was then filtered through a 200 µm nylon mesh and kept on ice. Within 10 minutes the homogenate was injected into the Oroboros Oxygraph-O2K chambers (Oroboros Instruments GmbH, Austria) containing MiR05 to measure a) mitochondrial respiration and hydrogen peroxide (H_2_O_2_) production under different respiratory states; b) oxygen affinity of complex IV and c) reverse electron transport capacity (RET). Remaining homogenate was frozen at -80°C and subsequently used for mitochondrial complex and citrate synthase activity assays.

Microrespirometry data were then normalised to citrate synthase activity ^35,36^.

### Mitochondrial oxygen consumption and ROS production

To examine mitochondrial respiration at different respiratory states, a substrate-uncoupler-inhibitor-titration (SUIT) protocol was employed as previously described^35,36^. H_2_O_2_ production was measured by adding Amplex® UltraRed (10 µM), 1 U.mL^-1^ horseradish peroxidase (HRP) and 5 U.mL^-1^ superoxide dismutase (SOD) into each chamber. Signal was calibrated by adding 40 µmol.L^-1^ H_2_O_2_ twice to achieve final concentrations of 0.1 and 0.2 μmol.L^-1^ in each chamber. Finally, complex I (CI) substrates: pyruvate (6.25 mmol.L^-1^), malate (2 mmol.L^-1^), and glutamate (10 mmol.L^-1^) were injected into each chamber prior to homogenate addition (approximately 100 – 150 µL; control: 0.087 – 0.43 mg.mL^-1^; HF: 0.052 – 0.19 mg.mL^-1^) to measure LEAK respiration with CI substrates (LEAK_CI_). Subsequently, ADP (5 mmol.L^-1^) and succinate (10 mmol.L^-1^) were added to measure oxidative phosphorylation with CI substrates (OXPHOS_CI_) and CII substrates (OXPHOS_CI+CII_), respectively. Mitochondrial uncoupler carbonyl cyanide p-trifluoro-methoxyphenyl hydrazone (FCCP) was added to uncouple mitochondria and therefore measure maximum electron transfer capacity (ETC_CI+CII_). FCCP was added stepwise to a final concentration of 0.5 - 1 µmol.L^-1^ – a point where there were no further increases in respiration. Subsequently, rotenone (0.15 µmol.L^-^) was added as a complex I inhibitor and maximum electron transport capacity with complex II substrates only (ETC_CII_) was recorded. Antimycin A (12.5 µmol.L^-^) was then added to block complex III and residual non-mitochondrial oxygen consumption (ROX) was recorded. Finally, N,N,N′,N′-tetramethyl-p-phenylenediamine (TMPD, 0.5 mmol.L^-1^) with ascorbate (2 mmol.L^-1^, added to prevent TMPD autooxidation) were injected into chambers to act as electron donor to complex IV and allow to measure maximum capacity of complex IV (CIV). Background non-mitochondrial oxygen consumption following the TMPD addition was measured at the end by using complex IV inhibitor, sodium azide (50 mmol.L^-1^). The respiratory control ratio (RCR; ratio of OXPHOS_CI_ and LEAK_CI_) and OXPHOS coupling-control ratio (1-(L/P), calculated as (1-(LEAK_CI_/OXPHOS_CI_)) were used to describe the ATP production efficiency.

### Mitochondrial oxygen affinity

In order to assess mitochondrial oxygen affinity, the respirometry chamber O_2_ concentration was adjusted to ∼ 50 µmol.L^-1^. CI substrates: pyruvate (6.25 mmol.L^-1^), malate (2 mmol.L^-1^), and glutamate (10 mmol.L^-1^) and homogenate (approximately 150 – 200 µL) were then added. Next, saturating concentrations of ADP (5 mmol.L^-1^) and succinate (10 mmol.L^-1^) were added to trigger OXPHOS_CI+CII_. O_2_ consumption rate was then monitored until both chamber O_2_ concentration and O_2_ consumption rate approached zero or fell below the lower limit of detection. For each sample, the same number of points were taken for analysis (approximately the last 4 minutes of the trace). The oxygen flux was corrected, plotted on a graph with and a nonlinear Michaelis–Menten regression curve was fitted. From the curve, the P_50_ (50% enzyme saturation) value was calculated for each sample in GraphPad Prism 8 software (GraphPad Software, Inc. USA).

### Reverse electron transport capacity

RET is the process by which electrons flow backwards to complex I, leading to the production of reactive oxygen species (ROS). SOD (5 U.mL^-1^), HRP (1 U.mL^-1^) and Amplex®UltraRed (10 µmol.L^-1^) and approximately 100 µL (0.08 – 0.15 mg.mL^-1^) of homogenate were added into each chamber. Baseline H_2_O_2_ production rate was recorded, and succinate (5 mmol.L^-1^) was added to promote RET to complex I. Signal was calibrated once by adding 40 µmol.L^-1^ H_2_O_2_ solution twice to achieve final concentration of 0.1 and 0.2 µmol.L^-1^ in each chamber.

### Western blot

The abundance of different mitochondrial proteins was measured by Western blot as previously described^37^. Briefly, CB tissue was homogenised in RIPA buffer containing protease and phosphatase inhibitor cocktail (1:100 dilution) and protein concentration was determined using a Bradford assay following manufacturer’s recommendations (*DC* protein assay kit, Bio-Rad, UK). Samples were run on a NuPAGE™ 4-12% Bis-Tris pre-cast polyacrylamide protein gel and transferred onto a nitrocellulose membrane (see Supplementary Table 1 for all details). The membrane was then incubated with a total protein stain and imaged before blocking with 5% milk. Membranes were first incubated with the primary antibodies overnight and then with the anti-rabbit secondary antibody. All blots were run in triplicate and averaged. The abundance of the target protein was normalised to the total protein stain or housekeeper (TOM20) and an internal control.

### Spectrophotometric analysis of enzymatic activities

Citrate synthase activity was determined by diluting homogenate samples in assay buffer containing 100 mmol.L^-1^ Tris, 300 µmol.L^-1^ acetyl coenzyme-A, 100 µmol.L^-1^ 5,5′-dithiobis-(2-nitrobenzoic acid)[DTNB], 0.1% Triton-X-100). The reaction was initiated by adding oxaloacetate (0.5 µmol.L^-1^). Following oxaloacetate addition, absorbance was measured repeatedly for 10 minutes at 412nm to determine V_max_. Values were then normalised to protein content in each homogenate sample (determined by Bradford method [Bio-Rad, UK], and corrected for the BSA content of the Mir05 buffer).

### Complex activity assays

Individual complex activity assays were performed as described previously^36,38^. Briefly, frozen tissue homogenates were diluted in phosphate buffer. Next, substrates for each individual complex were added (Supplementary Table 2). All samples were pre-incubated with complex substrates, and the fluorescence was read for approximately 2 minutes before the reaction was initiated by adding complex specific compounds. Fluorescence was then measured for another 10 minutes. The maximum rate of the reaction (V_max_) was calculated for each well. If normalised to protein concentration, the individual complex activity was calculated (Equation 1):

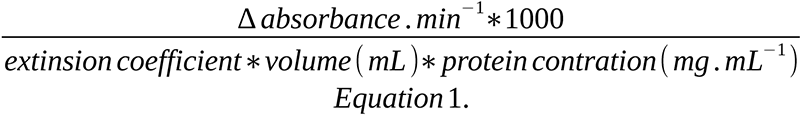

If normalised to citrate synthase, the individual complex activity was calculated (Equation 2):

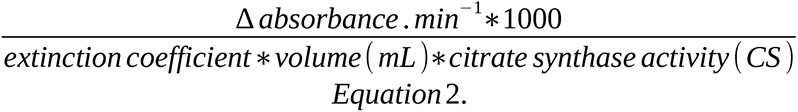

### Quantification of oxidative lipid damage

Approximately 30 – 40 mg of frozen tissue was placed 10x volumes homogenisation buffer containing: 10 mmol.L^-1^ phosphate buffer (pH = 7.2) and 200 µmol.L^-1^ butylated hydroxytoluene (used to prevent any artificial increases in oxidative stress due to the homogenisation process). Tissue was homogenised at 4°C and protein concentration in each sample was determined by using Bradford method. Tissue samples were added to a TBA-TCA-HCl buffer containing 0.375% thiobarbituric acid (TBA), 15% trichloroacetic acid (TCA) and 0.25 mol.L^-1^ hydrochloric acid (HCl) in 1:2 ratio. Samples were incubated for 1 hour at 95°C and then placed on ice for another 10 minutes. Product was then mixed with 1-butanol at 2:1 ratio and centrifuged at 15,000g for 3 minutes to separate malondialdehyde (MDA)-thiobarbituric acid adduct from the rest of solution. The top layer containing adduct (approximately 200 µL) was pipetted into a 96 well plate and absorbance was read at 532 nm. Throughout 1,1,3,3-tetraethoxypropane was used as a standard, which allowed for MDA levels calculation. MDA concentration was normalised to protein concentration and expressed as nmoles.mg^-1^ protein.

### Mitochondrial oxidative stress assay

CB tissue (∼ 50 – 90 mg) was homogenised in 3x volumes of ice-cold isolation buffer containing (in mmol.L^-1^): Tris hydrochloride (Tris-HCl; pH = 7.4),50; manganese chloride, 0.6; sodium citrate, 2; and butylated hydroxytoluene, 0.2. Following homogenisation, tissue homogenate was centrifuged at 600g for 10 minutes at 4°C, supernatant was kept. Assay buffer containing (in mmol.L^-1^): Tris-HCl (pH 7.4), 50; manganese chloride, 0.6; sodium citrate (pH 7.0), 5; 0.2 NADP^+^,0.2; 0.1% volume/volume Triton X-100, 0.4 U.mL^-1^ isocitrate dehydrogenase was prepared and warmed up to 30°C. 10 µL of each sample was pipetted into a 96-well plate and absorbance was read at 340 nm at regular intervals (with low shaking every 3 minutes) for 20 minutes at 30°C. The stable reading at the end of the protocol was taken for analysis. The aconitase activity was normalised to the protein content in each sample, which was measured by using a Bradford method.

### Statistics

All data analysis and graph preparation were performed using GraphPad Prism, version 8. Data are presented as mean ± standard deviation (SD). Each dataset was tested for normality of distribution by Shapiro-Wilk test. Where data was not normally distributed, values were transformed by using log(Y) and tested again ^39^. When data was normally distributed data, parametric t-tests were used as indicated in the figure legends. In cases when data was not normally distributed, nonparametric (Mann-Whitney) test was used (see figure legends).

Exact p values are displayed on each graph for p > 0.0001; a p value cut off of p < 0.05 was taken to infer statistical significance between groups. All data points are presented.

## Results

### Heart failure development

All animals implanted with a pacemaker developed subjective signs of HF (lethargy and breathlessness) on average after 58.1 ± 14 days (range 36 – 87 days) or tachypacing. Cardiac contractility was impaired showing a decrease in ejection fraction (67.9 ± 7.4 % pre-pace vs 26.5 ± 6.3 % HF, p <0.0001, n = 10; see ^32^ for a full characterisation of the model).

Furthermore, an increase in sympathetic activity was observed during the HF development (Supplementary Figure 1). However, neither a decrease in cardiac function nor increase in sympathetic control of HR is exclusively indicative of the CB dysfunction hence sheep underwent spirometry assessment to measure the hypoxic ventilatory response (HVR). Exposure to 11.5% FiO_2_ resulted in an increase in R_f_ pre-pace and in HF while V_T_ remained unchanged in response to hypoxia pre-pace but was increased in HF (Figure 1A-B). As a result, V_E_ was statistically increased in response to hypoxia both pre-pace (39.3 ± 31 % increase) and in HF (65.6 ± 47 % increase). Finally, the ΔV_E_ was higher by 96 ± 40 % in HF animals (Figure 1C) indicating that the CB signalling was indeed enhanced in the HF group. Additionally, animals with the highest ΔV_E_ on the last day of the protocol, developed HF faster than those with lower ΔV_E_ (Figure 1D). Overall, these data indicate that CB overactivation was present in HF sheep and was linked to a faster disease progression.

**Figure 1.**
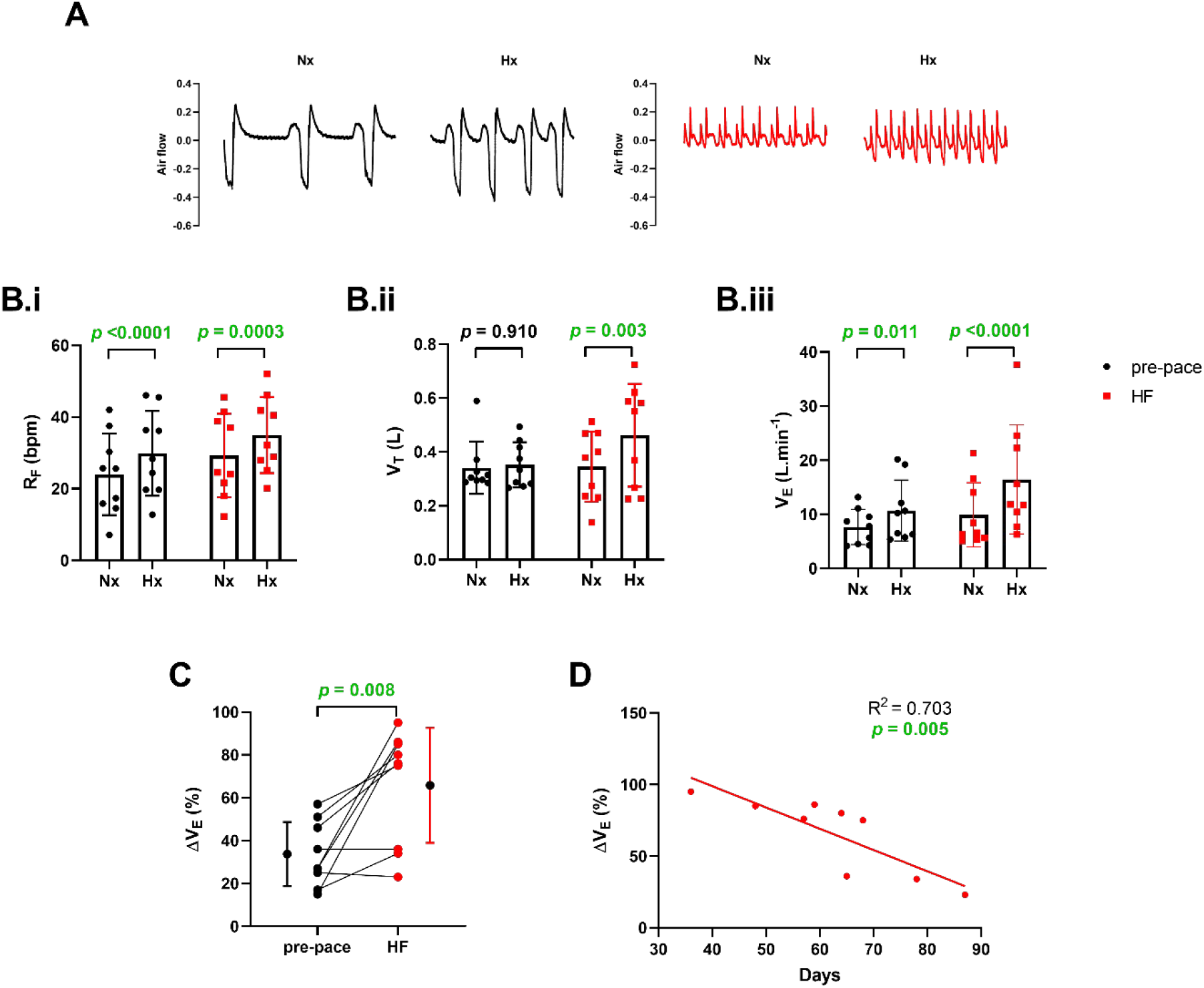
Characterisation of carotid body function before pacing protocol commenced (pre-pace) and in heart failure (HF). **A**. Representative 20 second traces of breathing under hyperoxic (Nx, 100% FiO_2_) and hypoxic (Hx, 11.5% FiO_2_) conditions at pre-pace and in heart failure. **B**. ventilatory variables recorded under hyperoxic (Nx 100% FiO_2_) and hypoxic (Hx, 11.5% FiO_2_) conditions: **i**) respiratory frequency (R_F_); **ii**) tidal volume (V_T_) and **iii**) minute ventilation (V_E_); **C**. percentage increase in V_E_ (ΔV_E_) in response to mild hypoxia pre-pace and in HF; **D**. relationship between number of days required to reach end of protocol (HF) and final ΔV_E_; N = 9 anaesthetised sheep in each group; paired t-test (B-C) and linear regression (D).

### Mitochondrial function in HF CB

The role of mitochondria in driving the changes in CB activity in HF has not been fully elucidated, however given their potential role in the oxygen-sensing cascade, their function was assessed by high resolution respirometry (example trace in Figure 2A). Here, we observed that LEAK_CI_ was increased, while OXPHOS_CI_ was lower in the HF group (Figure 2B-C). OXPHOS_CI+CII_, ETC_CI+CII_, ETC_CII_ and CIV remained unchanged in HF (Figure 2D-G). Change in LEAK_CI_ resulted in a lower RCR and OXPHOS coupling ratios (Figure 2H-I) indicating lower ATP production efficiency in the HF group. To further explore the effects of HF on CB mitochondria, mitochondrial complex activity and protein abundance were measured.

**Figure 2.**
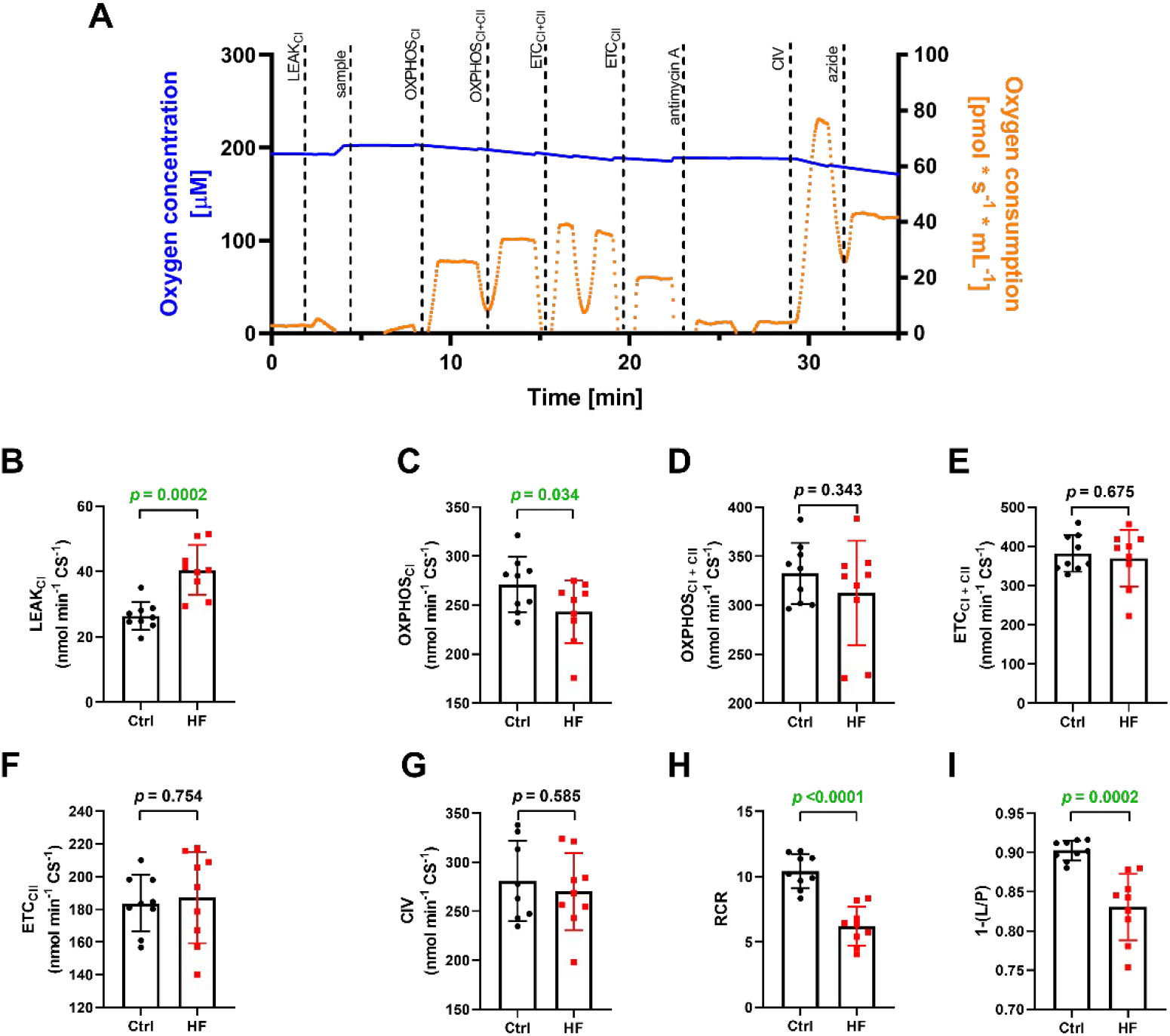
Carotid body (CB) mitochondria have higher LEAK, but a lower OXPHOS_CI_ respiration in heart failure (HF). **A**. Original traces of the SUIT protocol showing mitochondrial oxygen consumption under different respiratory states. Note that measurements were made when the trace was stable and injection artefacts excluded. **B**. LEAK respiration with complex I substrates (LEAK_CI_); **C**. oxidative phosphorylation with complex I substrates (OXPHOS_CI_); **D**. OXPHOS with complex I and complex II substrates (OXPHOS_CI+CII_); **E**. maximum electron transfer capacity with complex I and complex II substrates (ETC_CI+CII_); **F**. ETC with complex II substrates (ETC_CII_); **G**. maximum complex IV (CIV) activity; **H**. respiratory control ratio (RCR); **I**. LEAK/OXPHOS coupling-control ratio (1-(L/P)); n = 9 CBs in each group; two-way unpaired t-test (B, DC-IH), one-way unpaired t-test (C).

Firstly, we observed an increase in tyrosine hydroxylase (TH) and glial fibrillary acidic protein (GFAP) – markers of type I and type II cell populations, respectively (Supplementary Figure 2). Despite an increase in TH and GFAP, there were no changes in abundance of citrate synthase, TOM20 and OPA1 (three main mitochondrial markers) and complex I in HF (Supplementary Figure 3A-B & Figure 3A-B). The abundance of complexes II, III, IV and V was increased in the HF group when normalised to total protein (Figure 3C-F). However, when normalised to the housekeeper (TOM20), the increase in abundance was only observed for CIII and CIV (Supplementary Figure 3C-G). Finally, citrate synthase activity was increased in the HF group (Figure 4A) whilst individual complex enzyme activities remained unchanged when normalised to either citrate synthase activity or protein (Figure 4B-E & Supplementary Figure 4). Overall, these results indicate an increase in mitochondrial complex protein abundance with no changes in complex activity in HF.

**Figure 3.**
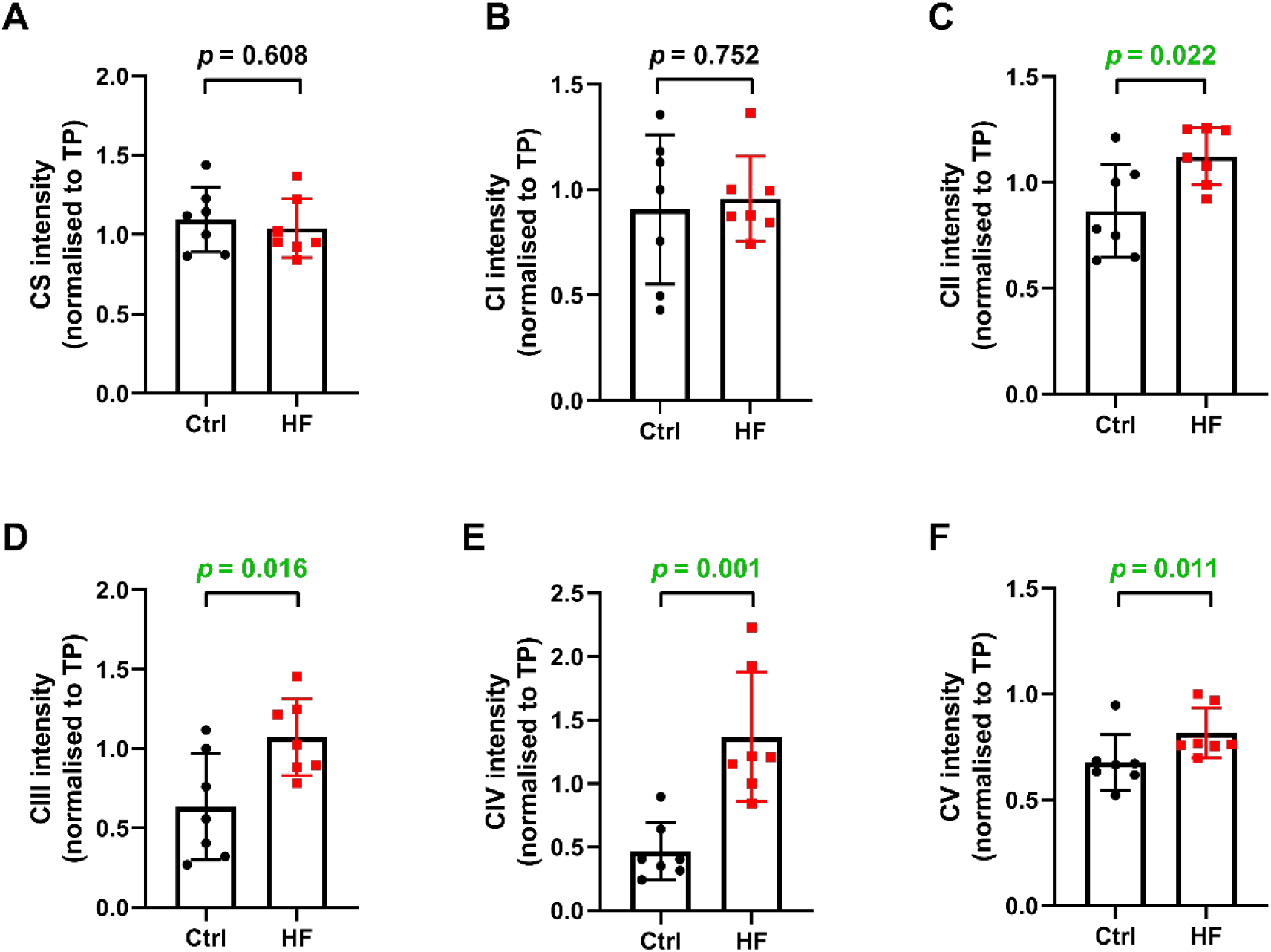
Abundance of CB complex II, complex III, complex IV and complex V is increased in heart failure (HF). Abundance of **A**. citrate synthase (CS); **B**. complex I (CI), **C**. complex II (CII), **D**. complex III (CIII), **E**. complex IV (CIV) and **F**. complex V (CV) normalised to total protein (TP); Ctrl – control; n = 7 CBs in each group, average of 3 technical repeats; unpaired t-test (A-D & E) and Mann-Whitney test (F).

**Figure 4.**
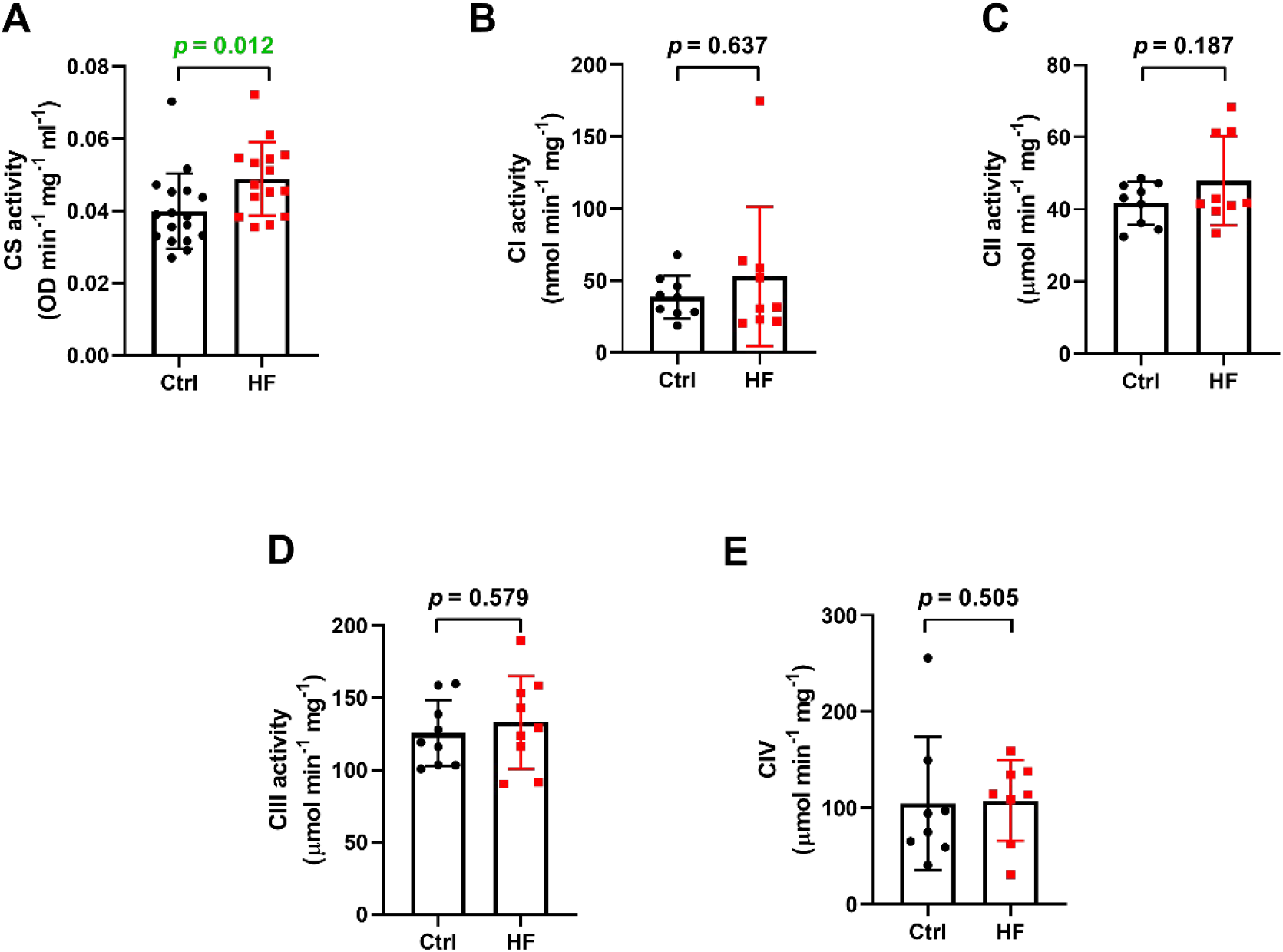
CB mitochondrial content, but not mitochondrial complex enzymatic activity is increased in heart failure (HF). Spectrophotometric analysis of enzymatic activity of **A**. citrate synthase (CS; Ctrl = 17 CBs, HF = 15 CBs), **B**. complex I (CI; n = 9 CBs in each group), **C**. complex II (CII; n = 9 CBs in each group), **D**. complex III (CIII; n = 9 CBs in each group) and **E**. complex IV (CIV; n = 8 CBs in each group) in tissue homogenates. Ctrl – control; parametric unpaired t-test (A-D) and Mann-Whitney test (E).

### Changes to the CB oxygen sensing cascade in HF

CB sensitivity can be modulated by a number of factors, including changes in mitochondrial oxygen affinity, specifically complex IV oxygen affinity. Equally any changes to the downstream mediators of the oxygen-sensing cascade could affect the oxygen-sensing cascade, hence we next measured oxygen affinity of complex IV and RET capacity, two key components of the mitochondria-to-membrane hypothesis. Example trace of oxygen affinity protocol is shown in Figure 5A. The oxygen affinity of complex IV was decreased in the HF group compared to the control (Figure 5D-E). As highlighted in the accompanying paper^36^, there was no biphasic response to falling levels of oxygen, instead the oxygen consumption remained stable through the recording (Figure 5B). At the same time, no changes to the RET capacity were found (Figure 5C&F). Together these results suggest that the oxygen sensitivity of complex IV was increased in HF CB and therefore could be contributing to an increased CB hypoxic sensitivity and increased hypoxic responsiveness *in vivo* in HF.

**Figure 5.**
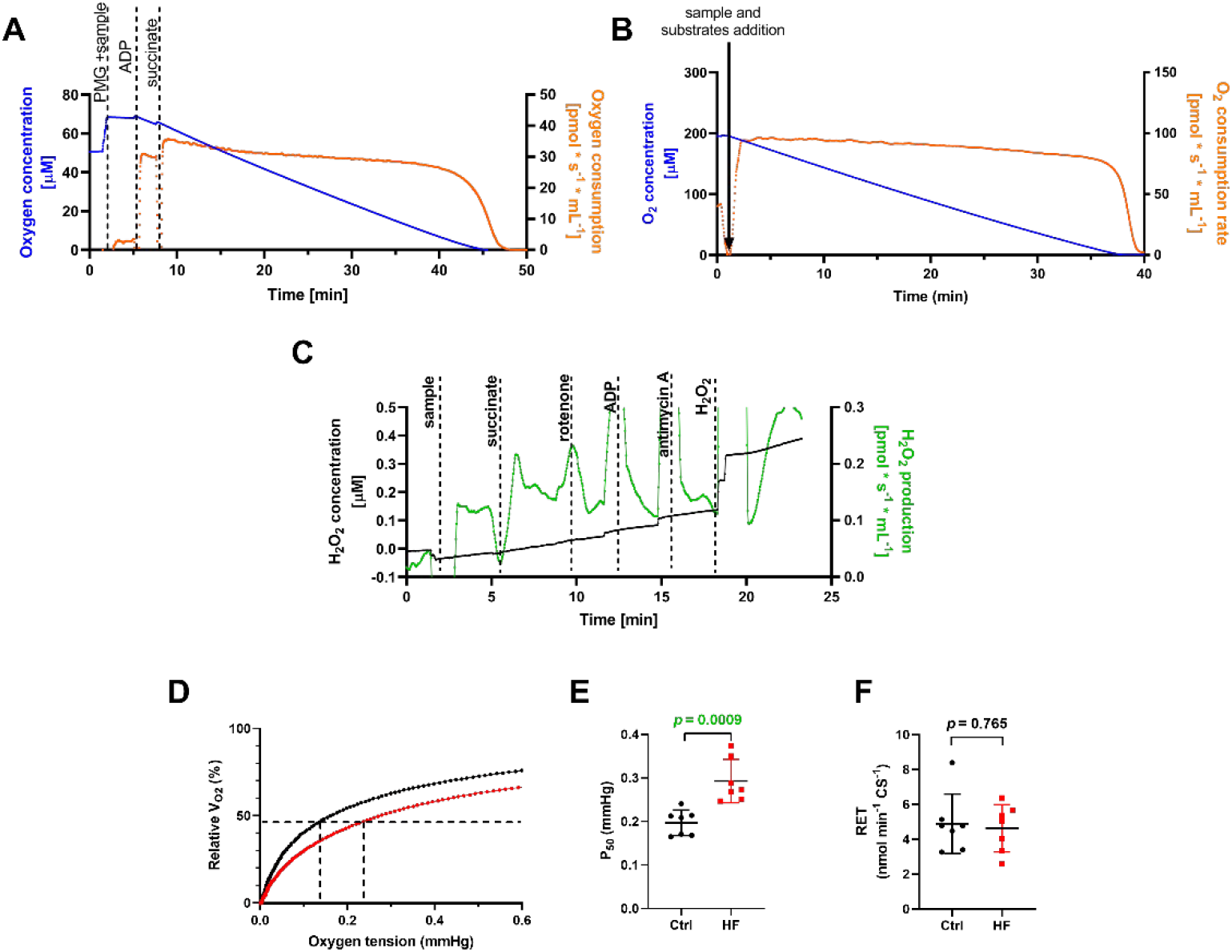
The cytochrome c oxidase oxygen affinity is decreased, while RET capacity remains unchanged in heart failure (HF) carotid body mitochondria. Representative traces of **A**. oxygen affinity protocol, **B**. oxygen consumption in carotid body (CB) homogenate sample when the chamber was fully oxygenated at the start of the protocol, **C**. reverse electron transport protocol showing changes in H_2_O_2_ production rate. **D**. relationship between relative oxygen consumption rate and oxygen tension; **E**. 50% enzyme saturation value (P_50_); H_2_O_2_ production rate at **F**. RET; E: n = 7 CBs in each group; F: Ctrl = 8 CBs; HF = 7 CBs, unpaired t-test; ADP - adenosine 5′-diphosphate, PMG – pyruvate, malate, glutamate.

### Oxidative stress in HF

Oxidative stress has been shown to have a role in driving CB overactivation in HF, hence we then measured cytosolic and mitochondrial oxidative stress using different approaches.

Firstly, we were able to show an increase lipid peroxidation in the HF group (Figure 6B) indicating a general increase in oxidative stress potentially driven by an increase in abundance of angiotensin II type 1 receptor and NADPH oxidase 2 (Figure 7). In order to specifically focus on mitochondrial oxidative stress, an aconitase assay was performed. Aconitase is one of the enzymes of Krebs cycle susceptible to ROS damage, namely aconitase activity decreases as H_2_O_2_ concentration increases (Supplementary Figure 5). In our study we show an increase in aconitase activity therefore suggesting a decrease in mitochondrial oxidative stress (Figure 6C). We then measured the mitochondrial H_2_O_2_ production by high resolution respirometry (Figure 6A). The mitochondrial respirometry data showed that ROS production was indeed lower during LEAK_CI_ respiration in HF (when normalised to oxygen flux, but not to citrate synthase; Figure 6D & Supplementary Figure 6A, respectively). No changes in ROS production under OXPHOS or ETC respiration were observed (Figure 6E-I & Supplementary Figure 6B-F). Overall, these results show that mitochondria may not have a role in driving oxidative stress in HF CB.

**Figure 6.**
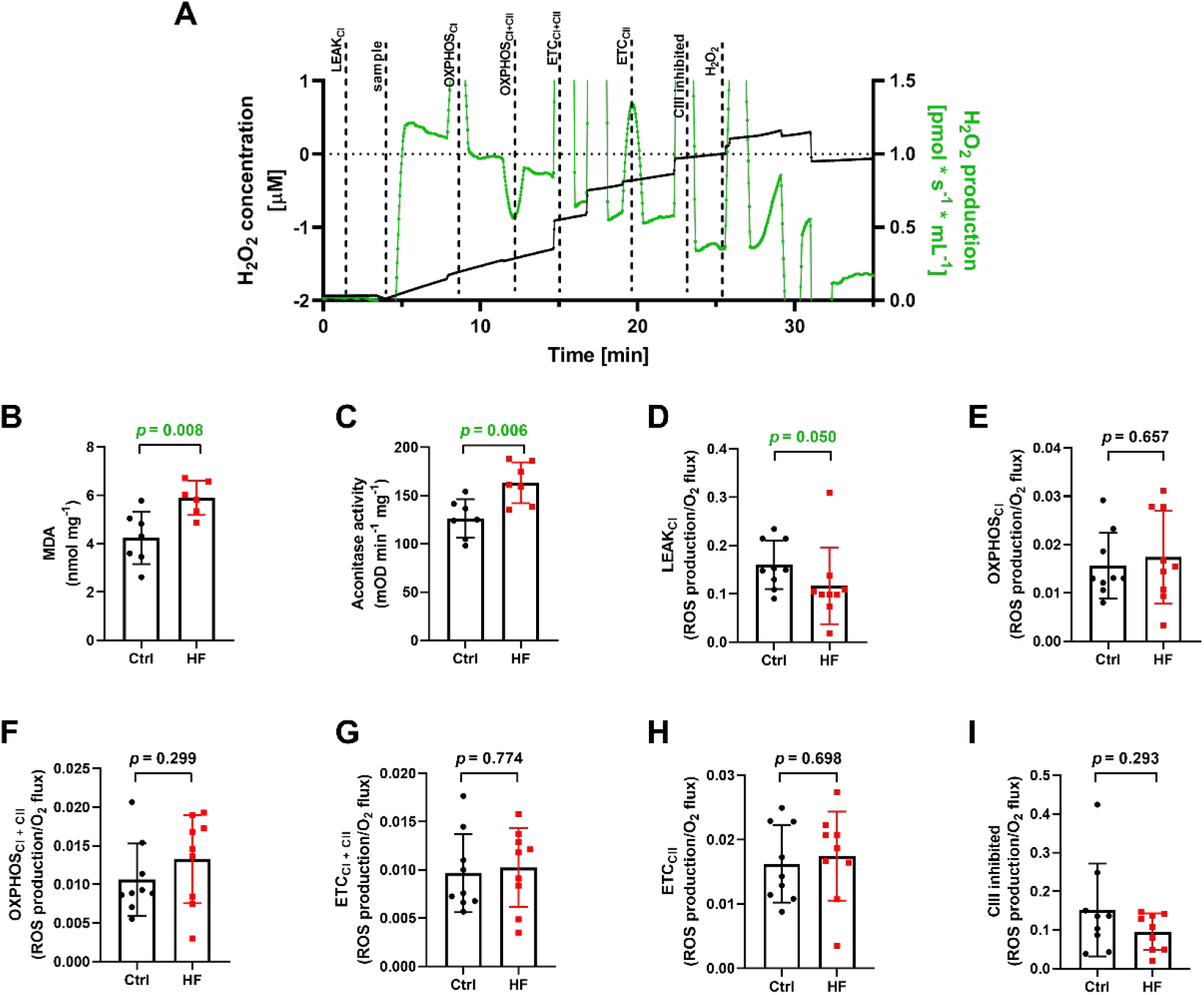
Both, CB global oxidative stress and aconitase activity are increased, while CB mitochondrial H_2_O_2_ production rate is decreased during LEAK respiration, but not other respiratory states in heart failure (HF). A. Original traces of the SUIT protocol showing H_2_O_2_ production measurements under different respiratory states. Prior to experimentation, Amplex UltraRed, horseradish peroxidase and superoxidase dismutase were added to measure the H_2_O_2_ production. Note that measurements were made when the trace was stable and injection artefacts excluded. B. malondialdehyde (MDA) levels; C. aconitase activity. Mitochondrial H_2_O_2_ production rate during D. LEAK respiration with complex I substrates (LEAK_CI_); E. oxidative phosphorylation with complex I substrates (OXPHOS_CI_); F. OXPHOS with complex I and complex II substrates (OXPHOS_CI+CII_); G. maximum electron transfer capacity with complex I and complex II substrates (ETC_CI+CII_); H. ETC with complex II substrates (ETC_CII_); I. after CIII inhibition; A: n = 7 CBs in each group; B: Ctrl = 7 CBs, HF = 6 CBs; C-H: n = 9 CBs in each group; parametric unpaired t-test (B-C; E-ID-H) and unpaired Mann-Whitney test (D).

**Figure 7.**
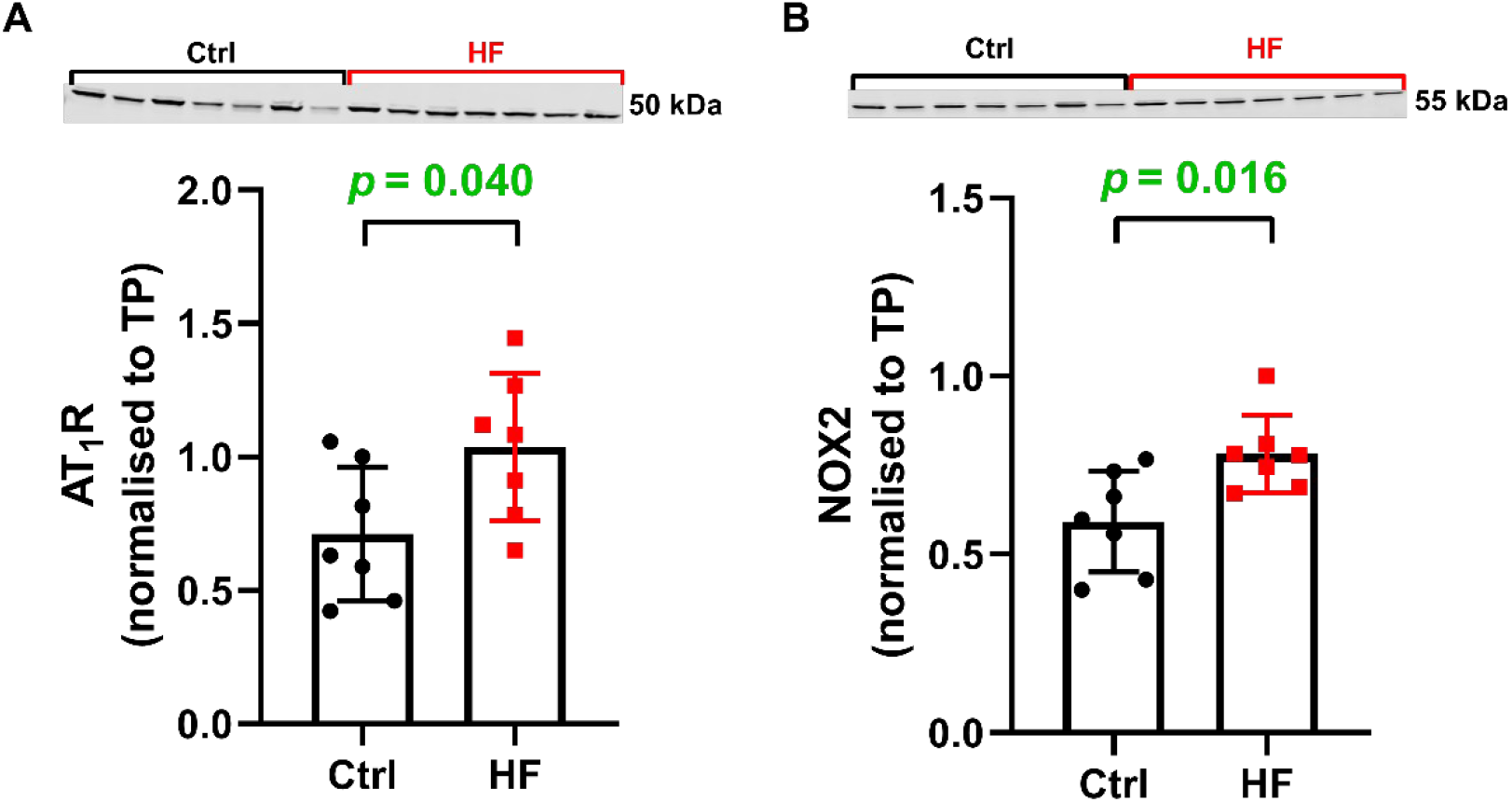
Abundance of A. angiotensin type I receptor (AT_1_R) and B. NADPH oxidase 2 (NOX2) are increased in the carotid body in heart failure (HF). Ctrl – control; TP – total protein, n = 7 CBs in each group, average of 3 technical repeats; unpaired t-test.

### Discussion

#### Main findings

The principal findings from this study are four-fold: I) we showed that hypoxic ventilatory response in elevated in HF and the magnitude of that increase is linked with a faster disease progression. II) The oxygen affinity of complex IV was lower suggesting an increased oxygen sensitivity which could be driving an increased HVR in the HF group. III) LEAK respiration was increased, which resulted in a decreased ATP production efficiency in HF. IV) Oxidative stress in HF CB was increased, however mitochondrial ROS production was lower suggesting that the increase in oxidative stress was more likely to be solely driven by cytosolic/extracellular enzymes. Overall, these results support the hypothesis that the CB mitochondrial complex IV contributes to driving an increased CB hypoxic sensitivity in HF. Nevertheless, the P_50_ value of complex IV remained low, indicating that the mitochondrial oxygen affinity was still high in the HF.

#### Oxygen sensitivity of the CB oxygen sensing cascade is increased in HF

CB hypoxic sensitivity could be driven by multiple factors either directly (changes to the components within the CB oxygen sensing cascade) or indirectly (via cytosolic enzymes/molecules that could affect the function of the sensor or ion channels involved in the chemotransduction cascade). Firstly, this study focused on assessing the oxygen affinity of complex IV. Despite the fact that our study performed on CB mitochondria showed that complex IV oxygen affinity is high and therefore contested previous findings^36^, any changes to complex IV oxygen affinity could still have a modulatory effect on the whole cell hypoxic sensitivity as mitochondria may still be an important part of the chemosensory pathway.

Here results showed that there was an increase in P_50_ suggesting the complex IV from HF CB mitochondria had lower oxygen affinity and therefore was more sensitive to any changes in the oxygen levels. Such change in the mitochondrial oxygen affinity could result in an enhanced response to hypoxia. Critically, the P_50_ values remain consistent with the role in oxygen sensing in HF, the actual values reported here are an order of magnitude lower than physiological oxygen levels within the mitochondria^40^.

One potential explanation for a decrease in oxygen affinity is a change in the subunit composition of complex IV. In particular, alterations in the expression of the COX4 isoforms (COX4i1 and COX4i2) have been shown to modulate the oxygen sensitivity of complex IV^41^. Specifically, increased incorporation of the COX4i2 isoform has been associated with enhanced oxygen sensitivity of complex IV^41,42^. The subunit composition of complex IV was not examined in the present study; however, investigating this could help determine whether changes in complex IV subunit composition are the primary mechanism responsible for the enhanced oxygen sensitivity observed in the carotid body during heart failure.

#### Carotid body mitochondrial function remains unchanged in HF

Mitochondria are important part of the cell metabolism and while it is widely reported in the literature that the mitochondrial oxidative capacity is decreased in failing hearts ^43,44^, there is no data on how CB mitochondrial function is affected by HF. Here, we show that there is an increase in LEAK_CI_ respiration suggesting that there is a higher level of proton slipping in the HF CB mitochondria. Such changes in LEAK respiration could be driven by several factors, including an increase in uncoupling proteins or a damage to mitochondrial membrane due to increased lipid peroxidation^45,46^. Increased LEAK_CI_ respiration can have two main outcomes: a) a decrease in protonmotive force, and b) protection against increase ROS production. Previous studies have shown that an increase in proton leak results in a decrease in protonmotive force and therefore a decrease in coupling efficiency^46,47^. Indeed, there was an approximately 40% fall in ATP production efficiency in the HF group compared to the control.

Mitochondrial function is regulated by a multitude of factors, including protonmotive force, as well as enzyme activity and protein abundance of the individual complexes. Previous studies on cardiac mitochondria showed that in HF, mitochondrial complex I, III and IV enzyme activity was decreased which then resulted in lower overall oxidative capacity^43^. Here, the abundance of citrate synthase and CI was unchanged, however CII, CIII, CIV and CV were increased in HF group, suggesting an increase in abundance of mitochondrial complexes. We observed that the citrate synthase activity was higher in the HF group, but no differences in enzyme activity of the individual complexes were detected. In comparison to the functional data that showed that the ATP production efficiency was decreased, enzymatic assays suggest that it was not driven due to a decrease in enzyme activity of each mitochondrial complex. Overall, these findings suggest that HF is associated with increased mitochondrial complex protein abundance without corresponding changes in complex activity, indicating that increased protein expression does not necessarily translate into enhanced functional capacity.

#### Carotid body oxidative stress is increased in HF

The presence of an increased oxidative stress in the CB tissue from HF models has been previously reported in the literature^48,49^. Later studies provided evidence that the increase in oxidative stress is driven by the increase in angiotensin II signalling and subsequent increase in NADPH oxidase 2 abundance and activity^48,50,51^. Furthermore, the antioxidant capacity (both mitochondrial and cytosolic SOD isoforms) within the CB was decreased^48,49,52^.

Together these data suggest that there is an increase in ROS production which is coupled with lack of/impaired ROS scavenging system in HF CB which in turn lowers the open probability of potassium channels^48,49,52^. One study also reported an increase in MitoSOX immunofluorescence in HF CB tissue which was localised to the type I cells^49^, suggesting mitochondria could be contributing to driving oxidative stress in HF. Here, we showed that while there was an increase in oxidative stress in CB tissue, mitochondrial ROS production remained the same HF. This was further supported by the fact that aconitase activity remained high in HF.

Aconitase activity is particularly susceptible to a rise in oxidative stress which suppresses its activity^53^. Here we showed that the aconitase activity was increased in HF CB suggesting no change or lower oxidative stress level in HF CB mitochondria. Combining that with increased LEAK_CI_ respiration, it is plausible that ROS production could remain unchanged or decrease as shown previously^47,54^. Indeed, an increase in LEAK_CI_ can decrease the protonmotive force via proton slipping, which results in ‘mild uncoupling‘, which ultimately leads to a lower ROS production ^54^. Lowering of the ROS production via the mild uncoupling was then proposed to be an adaptation process to prevent oxidative damage to the tissue under certain conditions (Brand’s hypothesis)^47^. Providing that the oxidative stress was detected in the CB tissue, it is therefore unlikely that mitochondria are contributing towards driving oxidative stress in HF. The increased levels of lipid peroxidation, angiotensin II type 1 receptor (AT₁R) and NADPH oxidase 2 suggest that oxidative stress in HF may be driven, at least in part, by enhanced angiotensin II signalling and subsequent activation of NADPH oxidase 2.

#### Limitations

All experiments were performed using tissue homogenates. Although these preparations allow reliable assessment of mitochondrial function, the presence of other cellular components and cell types, including type II cells, may have influenced the measured outcomes although the P_50_ we report here are similar to the ones we previously reported in isolated mitochondria^36^.

#### Conclusions

In summary, CB mitochondrial complex IV oxygen affinity was decreased in HF, suggesting that the CB mitochondria are more sensitive to oxygen changes. Such increase in oxygen sensitivity of complex IV could be a factor driving the increase in CB function in a HF setting. Additionally, while, there was an increase in oxidative stress in CB tissue from HF animals, mitochondria are not one of the contributing factors. On the contrary, mitochondrial ROS production may be decreased or unchanged due to ‘mild uncoupling’. Nevertheless, the increase in ROS production in HF CB tissue could still affect the ion channels and therefore result in an increased CB tonic activity. Overall, this is the first study to provide evidence that CB hypoxic sensitivity in HF could be enhanced at two points: firstly, in the mitochondria due to change in oxygen affinity of complex IV and secondly in the cytosol by an increase in oxidative stress. Oxidative stress can also have a role in enhancing CB tonic activity and therefore contributing towards sympathetic overdrive in HF patients.

### Sources of Funding

This work was supported by the British Heart Foundation (Grants: FS/19/60/34899; PG/23/11658; FS/CRTF/21/24140; PG/19/63/34601; PG/23/11642) and Medical Research Council (MR/Y003594/1).

### Author contribution statement

**Agnieszka Swiderska -** Writing – review & editing, Writing – original draft, Visualization, Validation, Methodology, Investigation, Formal analysis, Data curation, Conceptualization. **Mohammed Obeidat** – Methodology, Data curation, Investigation, Formal analysis, Writing – review & editing. **George W. P. Madders** – Methodology, Writing – review & editing. **Alice S. Whitley** – Methodology, Writing – review & editing. **Jagen C. Burke** – Methodology, Writing – review & editing. **Menglu Lin** - Methodology, Writing – review & editing. **Gina L.J. Galli:** Writing – review & editing, Writing – original draft, Supervision, Methodology, Conceptualization, Funding Acquisition, Investigation. **Andrew W. Trafford:** Writing – review & editing, Writing – original draft, Supervision, Project administration, Methodology, Investigation, Funding acquisition, Conceptualization.

## Ethics declaration

This study was conducted in accordance with the ARRIVE (Animal Research: Reporting of In Vivo Experiments) guidelines. This study was approved by the University of Manchester Animal Welfare and Ethical Review Board. (Approval No. PP3935083).

## Data Availability statement

Data associated with this project is available via a CC-BY Creative Commons Attribution licence through The University of Manchester FigShare repository available at (to be added at acceptance).

## Supplementary data

**Table 1.** Summary of Western blot running conditions and reagent sources.

| **Protein of interest** | **Amount of protein loaded (µg)** | **Running conditions** | **Primary antibody** | **1° Ab. Dilution** | **Secondary antibody** | **2° Ab. Dilution** |
|---|---|---|---|---|---|---|
| **NDUFS1 (complex I)** | 10 | MES, 200 V, 50 minutes | ProteinTech, 12444-1-AP, Rabbit polyclonal | 1:3000 | IRDye<sup>®</sup> 800CW Donkey anti-Rabbit IgG Secondary Antibody (LI-COR Biosciences, UK) | 1:20 000 |
| **SDHA (complex II)** | 10 | MES, 200 V, 50 minutes | ProteinTech, 14865-1-AP, Rabbit polyclonal | 1:2000 | IRDye<sup>®</sup> 800CW Donkey anti-Rabbit IgG Secondary Antibody (LI-COR Biosciences, UK) | 1:20 000 |
| **UQCRC2 (complex III)** | 10 | MES, 200 V, 50 minutes | ProteinTech, 14742-1-AP, Rabbit polyclonal | 1:4000 | IRDye<sup>®</sup> 800CW Donkey anti-Rabbit IgG Secondary Antibody (LI-COR Biosciences, UK) | 1:20 000 |
| **MTCO1 (complex IV)** | 10 | MES, 200 V, 50 minutes | Abcam, ab14705, Mouse monoclonal | 1:2000 | IRDye<sup>®</sup> 800CW Donkey anti-Mouse IgG Secondary Antibody (LI-COR Biosciences, UK) | 1:20 000 |
| **ATP5a (Complex V)** | 10 | MES, 200 V, 50 minutes | Abcam, ab14748, Mouse monoclonal | 1:1000 | IRDye<sup>®</sup> 800CW Donkey anti-Mouse IgG Secondary Antibody (LI-COR Biosciences, UK) | 1:20 000 |
| **CS** | 10 | MES, 200V, 50 minutes | ProteinTech, 16131-1-AP, Rabbit polyclonal | 1:2000 | IRDye<sup>®</sup> 800CW Donkey anti-Rabbit IgG Secondary Antibody (LI-COR Biosciences, UK) | 1:20 000 |
| **TOM20** | 10 | MES, 200 V, 50 minutes | ProteinTech, 11802-1-AP, Rabbit polyclonal | 1:1000 | IRDye<sup>®</sup> 800CW Donkey anti-Rabbit IgG Secondary Antibody (LI-COR Biosciences, UK) | 1:20 000 |
| **OPA1** | 10 | MES, 200V, 50 minutes | ProteinTech, 27733-1-AP, Rabbit polyclonal | 1:4000 | IRDye<sup>®</sup> 800CW Donkey anti-Rabbit IgG Secondary Antibody (LI-COR Biosciences, UK) | 1:20 000 |
| **AT<sub>1</sub>R** | 10 | MOPS, 150 V, 90 minutes | ProteinTech, 25343-1-AP, Rabbit polyclonal | 1:500 | IRDye<sup>®</sup> 800CW Donkey anti-Rabbit IgG Secondary Antibody (LI-COR Biosciences, UK) | 1:20 000 |
| **GFAP** | 10 | MOPS, 150 V, 90 minutes | Abcam, ab7260, Rabbit polyclonal | 1:10 000 | IRDye<sup>®</sup> 800CW Donkey anti-Rabbit IgG Secondary Antibody (LI-COR Biosciences, UK) | 1:20 000 |
| **NOX2** | 10 | MOPS, 150 V, 90 minutes | ProteinTech, 19013-1-AP, Rabbit polyclonal | 1:500 | IRDye<sup>®</sup> 800CW Donkey anti-Rabbit IgG Secondary Antibody (LI-COR Biosciences, UK) | 1:20 000 |
| **TH** | 10 | MOPS, 150 V, 90 minutes | Abcam, ab112, Rabbit polyclonal | 1:750 | IRDye<sup>®</sup> 800CW Donkey anti-Rabbit IgG Secondary Antibody (LI-COR Biosciences, UK) | 1:20 000 |

**Table 2.**
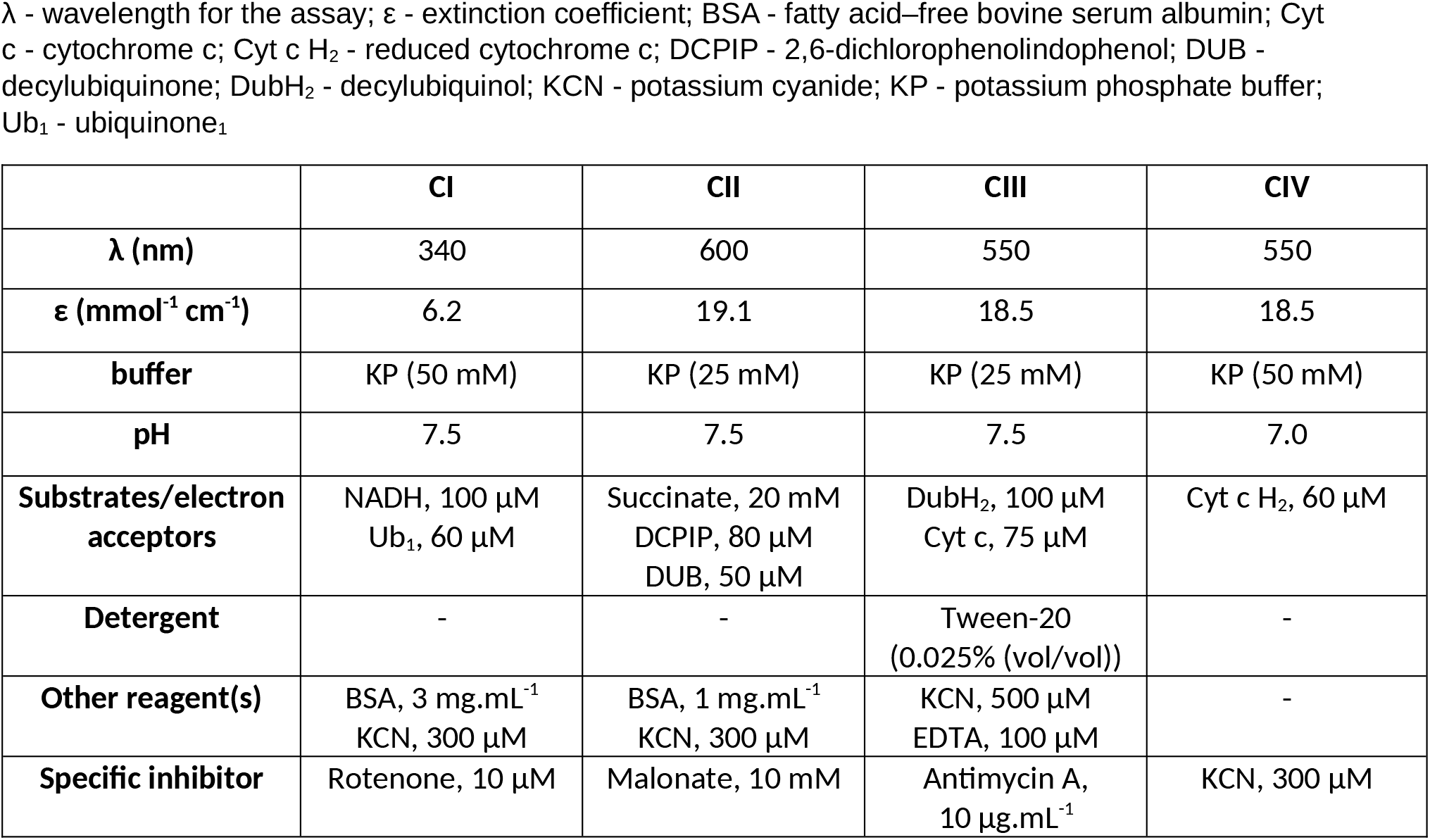
Conditions for mitochondrial complex activity assays.

|  | **CI** | **CII** | **CIII** | **CIV** |
|---|---|---|---|---|
| **λ (nm)** | 340 | 600 | 550 | 550 |
| **ε (mmol<sup>-1</sup> cm<sup>-1</sup>)** | 6.2 | 19.1 | 18.5 | 18.5 |
| **buffer** | KP (50 mM) | KP (25 mM) | KP (25 mM) | KP (50 mM) |
| **pH** | 7.5 | 7.5 | 7.5 | 7.0 |
| **Substrates/electron acceptors** | NADH, 100 µM<br>Ub<sub>1</sub>, 60 µM | Succinate, 20 mM<br>DCPIP, 80 µM<br>DUB, 50 µM | DubH<sub>2</sub>, 100 µM<br>Cyt c, 75 µM | Cyt c H<sub>2</sub>, 60 µM |
| **Detergent** | - | - | Tween-20 (0.025% (vol/vol)) | - |
| **Other reagent(s)** | BSA, 3 mg.mL<sup>-1</sup><br>KCN, 300 µM | BSA, 1 mg.mL<sup>-1</sup><br>KCN, 300 µM | KCN, 500 µM<br>EDTA, 100 µM | - |
| **Specific inhibitor** | Rotenone, 10 µM | Malonate, 10 mM | Antimycin A, 10 µg.mL<sup>-1</sup> | KCN, 300 µM |

**Supplementary Figure 1.**
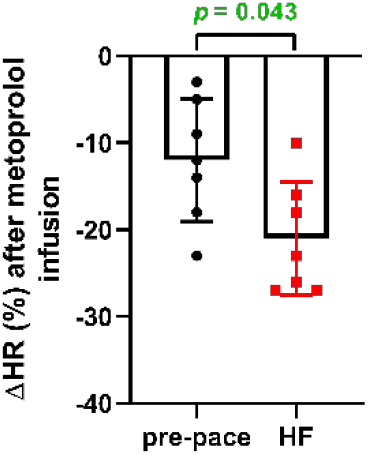
Heart rate responses to metoprolol pre-pacing and final week of heart failure (HF). Percent change in HR after metoprolol infusion at pre-pace and in HF, n = 7 conscious sheep in each group, parametric paired t-test.

**Supplementary Figure 2.**
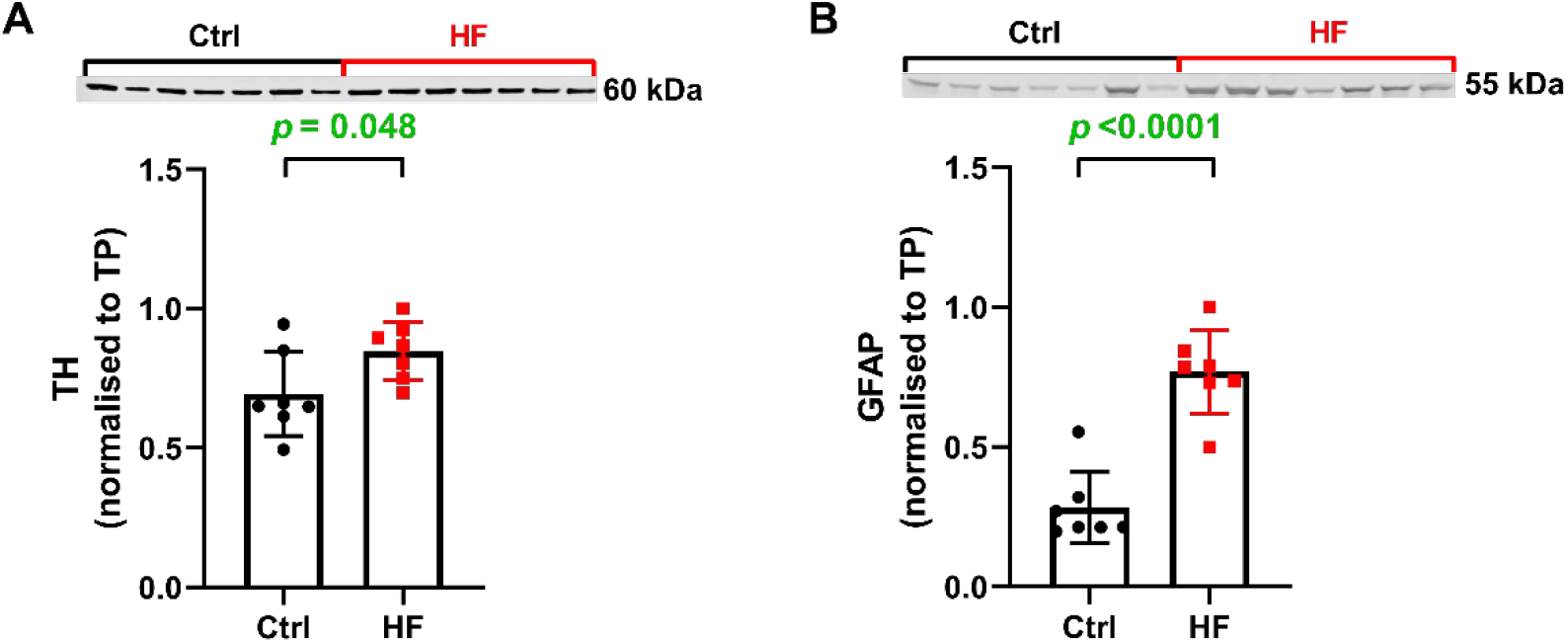
The abundance of A. tyrosine hydroxylase (TH) and B. glial fibrillary acidic protein (GFAP) are increased in heart failure (HF) carotid body. Ctrl – control; TP – total protein, n = 7 CBs in each group, average of 3 technical repeats, unpaired t-test.

**Supplementary Figure 3.**
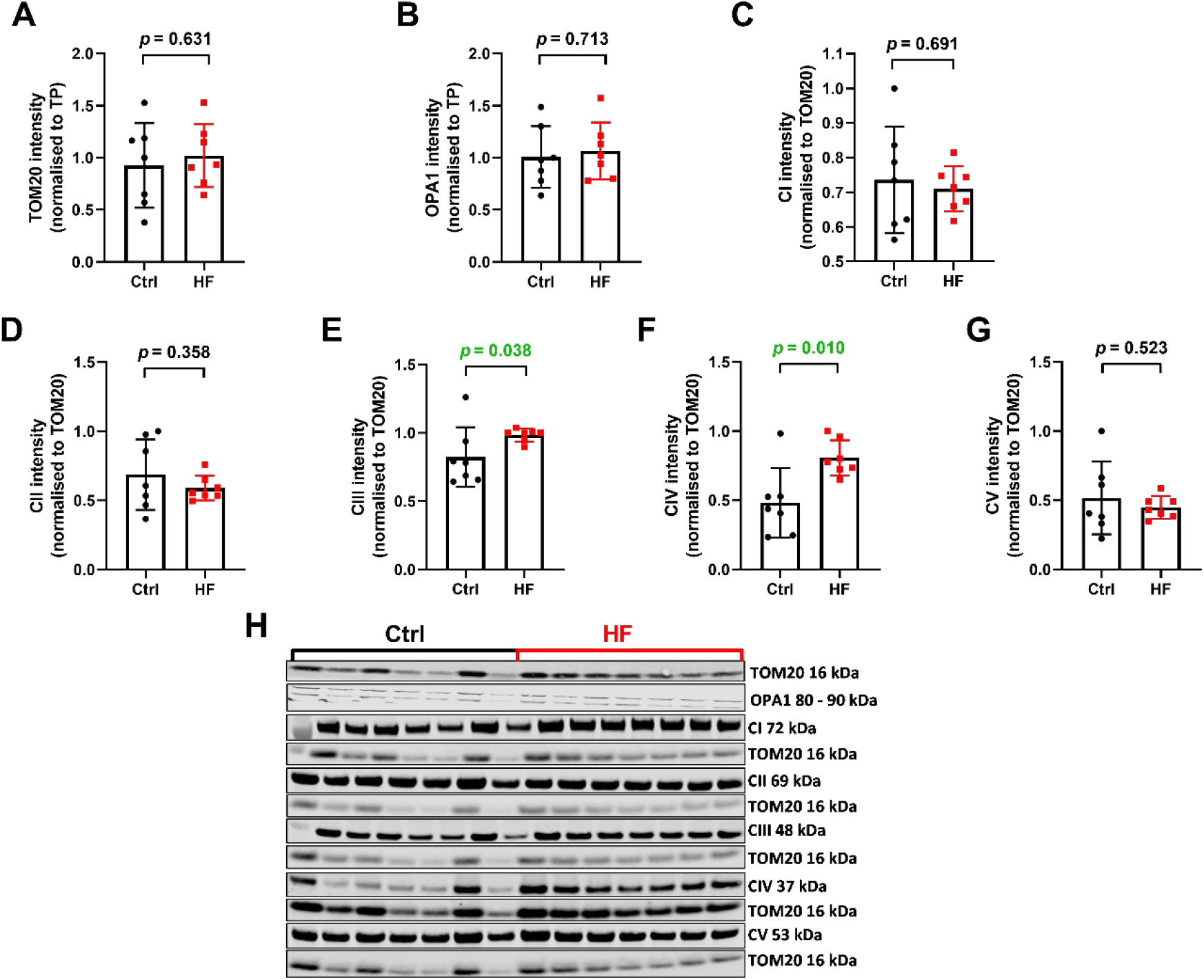
Complex III and complex IV abundance per TOM20 protein was increased in heart failure (HF) carotid body. Abundance of **A**. TOM20, **B**. OPA1, **C**. complex I (CI), **D**. complex II (CII), **E**. complex III (CIII), **F**. complex IV (CIV) and **G**. complex V (CV); **H**. representative western blot images, n = 7 CBs in each group, average of 3 technical repeats; unpaired t-test (A-D & F-G) and Mann Whitney test (E).

**Supplementary Figure 4.**
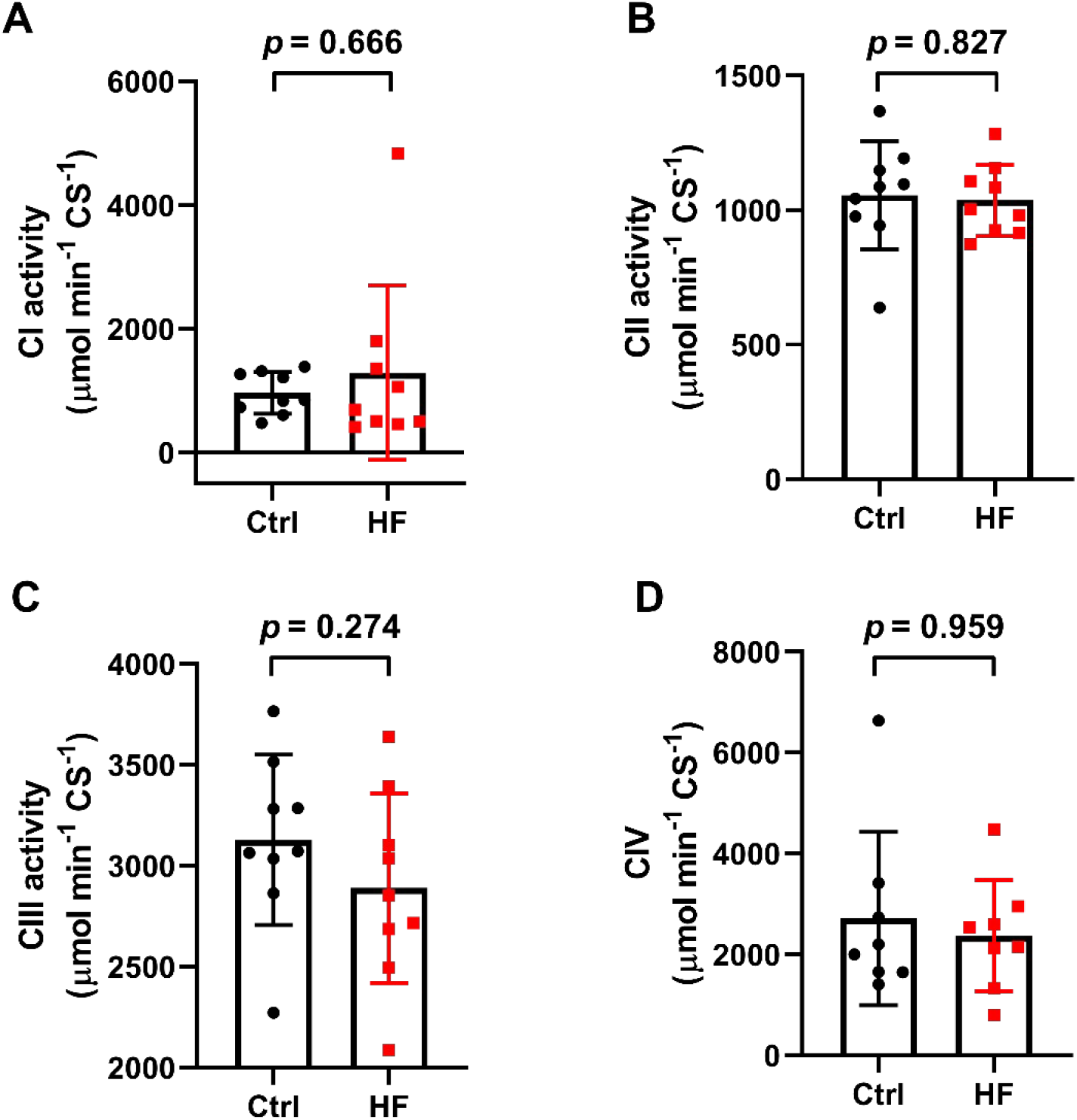
Enzymatic activity of individual mitochondrial complexes in heart failure (HF) remains the same when normalised to citrate synthase (CS). Enzymatic activity of **A**. complex I (CI; n = 9 CBs in each group), **B**. complex II (CII; n = 9 CBs in each group), **C**. complex III (CIII; n = 9 CBs in each group) and **D**. complex IV (CIV; n = 8 CBs in each group) per CS activity; Mann-Whitney test (A & D) and unpaired t-test (B-C).

**Supplementary Figure 5.**
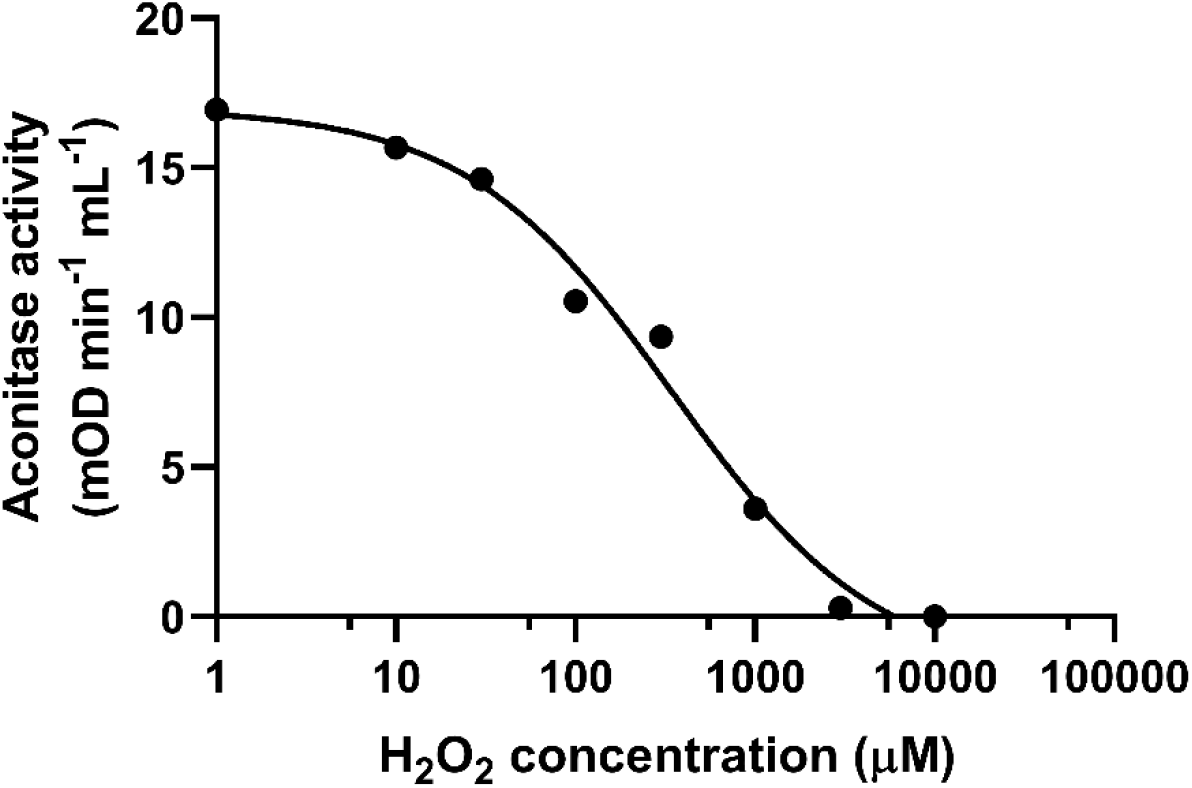
The relationship between the aconitase activity and increasing concentrations of hydrogen peroxide (H_2_O_2_).

**Supplementary Figure 6.**
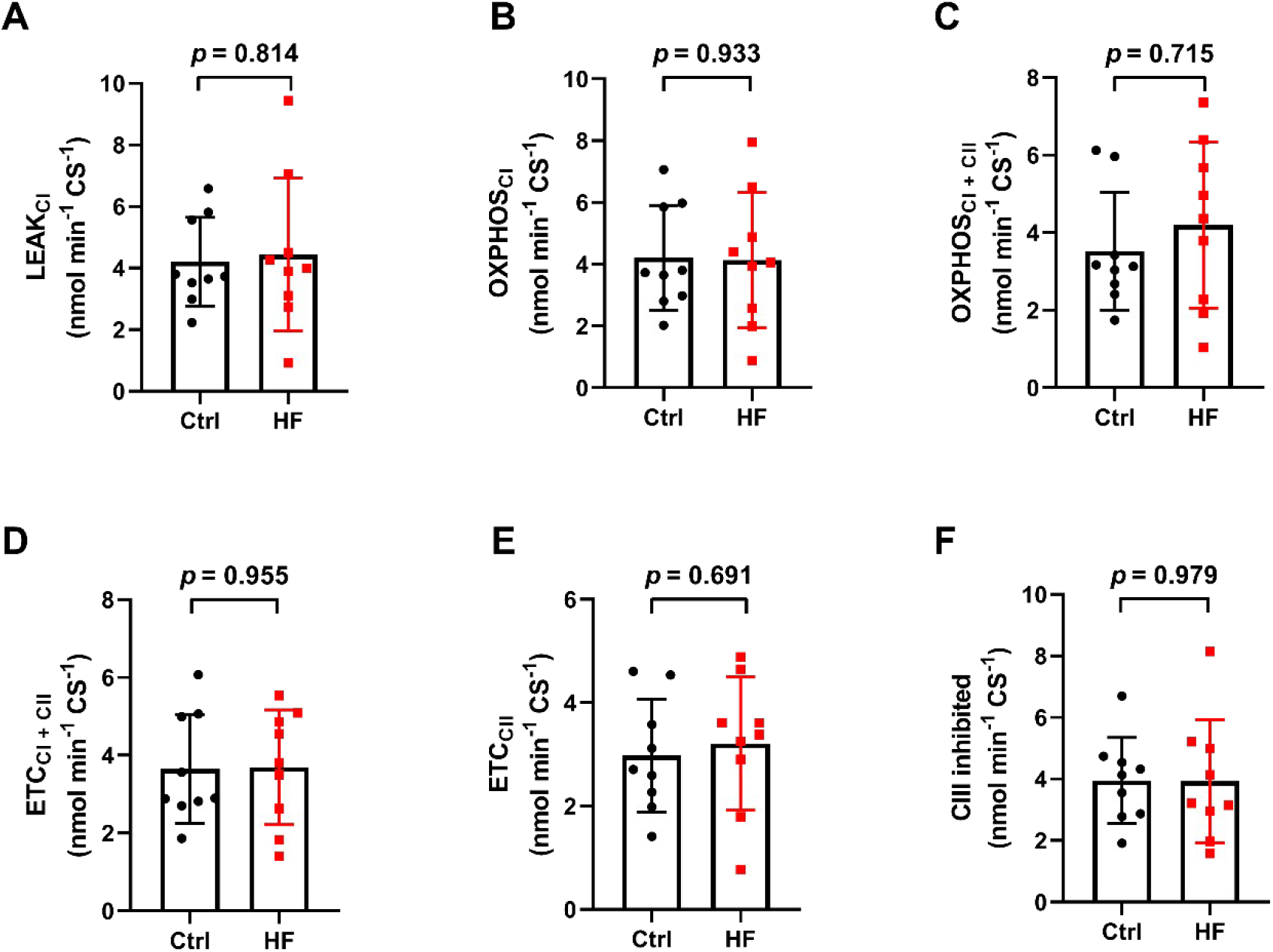
H_2_O2 production rate was unchanged in heart failure (HF) carotid body mitochondria when normalised to citrate synthase (CS) activity. H_2_O_2_ production rate during **A**. LEAK respiration with complex I substrates (LEAK_CI_); **B**. oxidative phosphorylation with complex I substrates (OXPHOS_CI_); **C**. OXPHOS with complex I and complex II substrates (OXPHOS_CI+CII_); **D**. maximum electron transfer capacity with complex I and complex II substrates (MAX_CI+CII_); **E**. MAX with complex II substrates (MAX_CII_); **F**. after CIII inhibition; n = 9 CBs in each group; unpaired t-test.

